# Transcriptional regulatory mechanisms of fibrosis development in mouse lung tissue exposed to carbon nanotubes

**DOI:** 10.1101/363762

**Authors:** Vadim Zhernovkov, Tapesh Santra, Hilary Cassidy, Oleksii Rukhlenko, David Matallanas, Aleksandar Krstic, Walter Kolch, Vladimir Lobaskin, Boris N. Kholodenko

## Abstract

**Background:** Carbon nanotubes (CNTs) usage has rapidly increased in the last few decades due to their unique properties, exploited in various industrial and commercial products. Certain types of CNTs cause adverse health effects, including chronic inflammation and fibrosis. Despite the large number of *in vitro* and *in vivo* studies evaluating these effects, many important questions remain unanswered due to a lack of mechanistic understanding of how CNTs induce cellular stress responses. In order to predict CNT toxicity, it is important to understand which transcriptional programs are specifically activated in response to CNTs, and what similarities and differences exist in relation to other toxic inducers exerting similar adverse effects.

**Results:** A systems biology approach was applied to reveal complex interactions at the molecular level in mouse lung tissue in response to different fibrosis inducers: two types of multi-walled CNTs, NM-401 and NRCWE-26, and bleomycin (BLM). Based on mRNA gene expression profiles, we inferred gene regulatory networks (GRNs) to capture functional hierarchical regulatory structures between genes and their regulators. We found that activities of the transcription factors (TFs) *Myc, Arid5a* and *Mxd1* were associated with the regulation of cytokine transcription in response to CNTs, while in response to BLM treatment, *Myc* was associated with p53 signaling. TF *Litaf* was identified as the essential regulator for noncanonical signaling of TLR2/4 driven by CNTs. Despite the different nature of the lung injury caused by CNTs and BLM, we identified common stress response modules, that included DNA damage (TFs: *E2f8, E2f1, Foxm1)*, M1/M2 macrophage polarization (TF: Mafb), Interferon response (TFs: *Irf7, Stat2* and *Irf9)* for all agents.

**Conclusions:** These results suggest that the reconstruction and analysis of TF-centric gene interaction networks can reveal key targets and regulators of cellular stress responses to toxic agents.

## Background

Carbon nanotubes (CNTs) have gained increased attention in the last 30 years due to their unique properties, exploited in various industrial and commercial products. CNTs are widely used in electronics, energy storage, composite materials, water filters, and biomedical applications, such as DNA/proteins biochip and drug delivery [1]. However, inhalation of certain CNTs causes adverse health effects, such as chronic inflammation, pulmonary fibrosis, carcinogenesis and other undesirable effects [2].

Accumulation of toxic CNTs in the lungs activates acute inflammatory responses. It initiates the recruitment of inflammatory cells including macrophages, lymphocytes and neutrophils and induces the secretion of proinflammatory cytokines, chemokines and growth factors. Initial response to nanoparticle inhalation also includes excessive extracellular matrix (ECM) remodeling and differentiation of fibroblasts into myofibroblasts [3]. This initial response then progresses to a chronic phase, which is characterized by decreased inflammatory processes, increased deposition of ECM proteins and myofibroblasts, and granuloma formation. Myofibroblasts are known to modulate the fibrosis progression by accumulating collagen fibers and other ECM proteins [4]. Other toxicological effects linked to CNT exposure include DNA genotoxicity and tumorigenesis [2]. CNTs are known to indirectly cause DNA damage, including DNA strand breaks and chromosomal aberrations, by inducing ROS generation or suppressing intracellular antioxidants [5, 6]. CNTs can also activate oxidative stress by directly interacting with the cell membranes, resulting in the increased concentrations of lipid peroxidation products. Frustrated phagocytosis is yet another mechanism of ROS generation, which occurs when phagocytes cannot engulf CNTs due to their rigidity and physical size [5, 6].

Many signaling pathways are involved in shaping biological responses to CNT exposure, with TGFβ_1_ signaling being one of the most important. The latent form of TGFβ_1_ is abundant in the ECM of healthy lung tissues. When activated, TGFβ_1_ triggers ECM remodeling by stimulating the secretion of pro-fibrotic proteins [7]. Besides its essential contribution to collagen deposition in ECM, TGFβ_1_ also plays a substantial role in the regulation of inflammation, macrophage recruitment, and the initiation of epithelial-mesenchymal transition [8]. In addition to TGFβ_1_ signaling, other signaling pathways, such as the p53 DNA damage response and NF-κB inflammatory pathways are also implicated in the CNT-induced fibrosis-related effects.

The biological response to CNT exposure has many similarities with the responses observed for bleomycin (BLM) treatment, that is widely used as a classic model of inducing lung fibrosis [9]. BLM is a chemotherapeutic drug that causes DNA strand breaks *via* oxidative mechanisms and can induce lung fibrosis as a severe side effect in human patients. A single BLM dose initiates an acute inflammatory phase in lung tissue characterized by infiltration of immune cells, release of pro-inflammatory cytokines, and the increased presence of myofibroblasts [10]. In rodents the fibrotic phase is initiated seven days after BLM instillation with increased expression of pro-fibrotic cytokines and increased fibroblast proliferation and collagen accumulation [10]. Elevated levels of the active form of TGFβ_1_ are detected in both acute and fibrotic stages. However, there are differences in endpoint effects of BLM and CNTs. For CNT exposure, genotoxic and pro-fibrotic responses together with immunomodulation components prevail, leading to chronic inflammation, fibrosis and possibly cancer [11].

A large number of studies have previously investigated the toxic effects of CNTs on lungs. These studies identified the genes and proteins which are influenced by various types of CNTs, as well as the pathways and biological process, in which these genes participated [12, 13]. However, important questions remain unanswered due to a lack of mechanistic understanding of how CNTs influence gene expression in lung cells. In order to predict CNT toxicity, it is important to understand which transcriptional programs are specifically activated in response to CNTs, and what similarities and differences exist compared to classic fibrosis inducers, such as BLM. To better understand and then prevent CNT induced fibrosis, we need to know whether all CNTs initiate fibrosis related processes *via* a shared mechanism or through unique mechanisms specific to their physico-chemical characteristics. Here, we apply a systems biology approach to reveal complex interactions at the molecular level in mouse lung tissue in response to different fibrosis inducers: two types of multi walled carbon nanotubes (MWCNTs) NM-401 and NRCWE-26 (that have different physico-chemical characteristics), and bleomycin (BLM). We inferred gene regulatory networks (GRNs) to capture functional hierarchical regulatory patterns connecting genes and their regulators. Using reverse engineering techniques and gene expression profiles from mRNA high-throughput experiments, we identified both gene regulators and their targets, which have key roles in pro-fibrotic progression for both MWCNTs and bleomycin-induced responses. These genes orchestrate specific transcriptional responses to MWCNT and bleomycin instillation. This integrative framework can also be applied to investigations of gene regulatory programs activated in response to different type of toxic agents.

## Results

To uncover the mechanisms of the pulmonary response to CNT and BLM exposure, transcriptomics data of previously published microarray experiments for MWCNTs NM-401 and NRCWE-26 [6] and BLM [9] were assessed. The gene expression profiles for two types of multi-walled carbon nanotubes (GSE55286), NM-401 (long and stiff) and NRCWE-26 (short and entangled), and BLM (GSE40151) were obtained from the Gene Expression Omnibus Database (https://www.ncbi.nlm.nih.gov/geo/). Data for CNTs and BLM were generated using Agilent SurePrint G3 Mouse GE 8×60K Microarrays and Affymetrix Mouse 430 2.0 arrays, respectively, with RNA isolated from mouse lung tissues. CNTs were instilled at three different doses of (18, 54, 162 μg), and gene expression was determined at three timepoints (1, 3, 28 days) post-instillation. BLM was administered in one dose (2U/kg body weight), and the lung tissue was harvested at 7 post-instillation timepoints (1, 2, 7, 14, 21, 28, 35 days). Both types of experiments were conducted with vehicle controls for each timepoint.

An overview schematic of our analysis pipeline is represented in Figure 1. First, using the reverse engineering algorithms and the transcriptomics data, we inferred a gene regulatory network for each agent. The gene interactions were identified for a combined list of DEGs derived from all experiments. Next, we identified network modules using the GLay clustering algorithm [14]. For each module, we performed pathway enrichment analysis to identify biological functions of genes in the module. To reveal TFs and signaling pathways directly associated with fibrosis development, we next analyzed subnetworks based on fibrosis markers and their direct regulators extracted from the whole networks.

**Figure 1.**
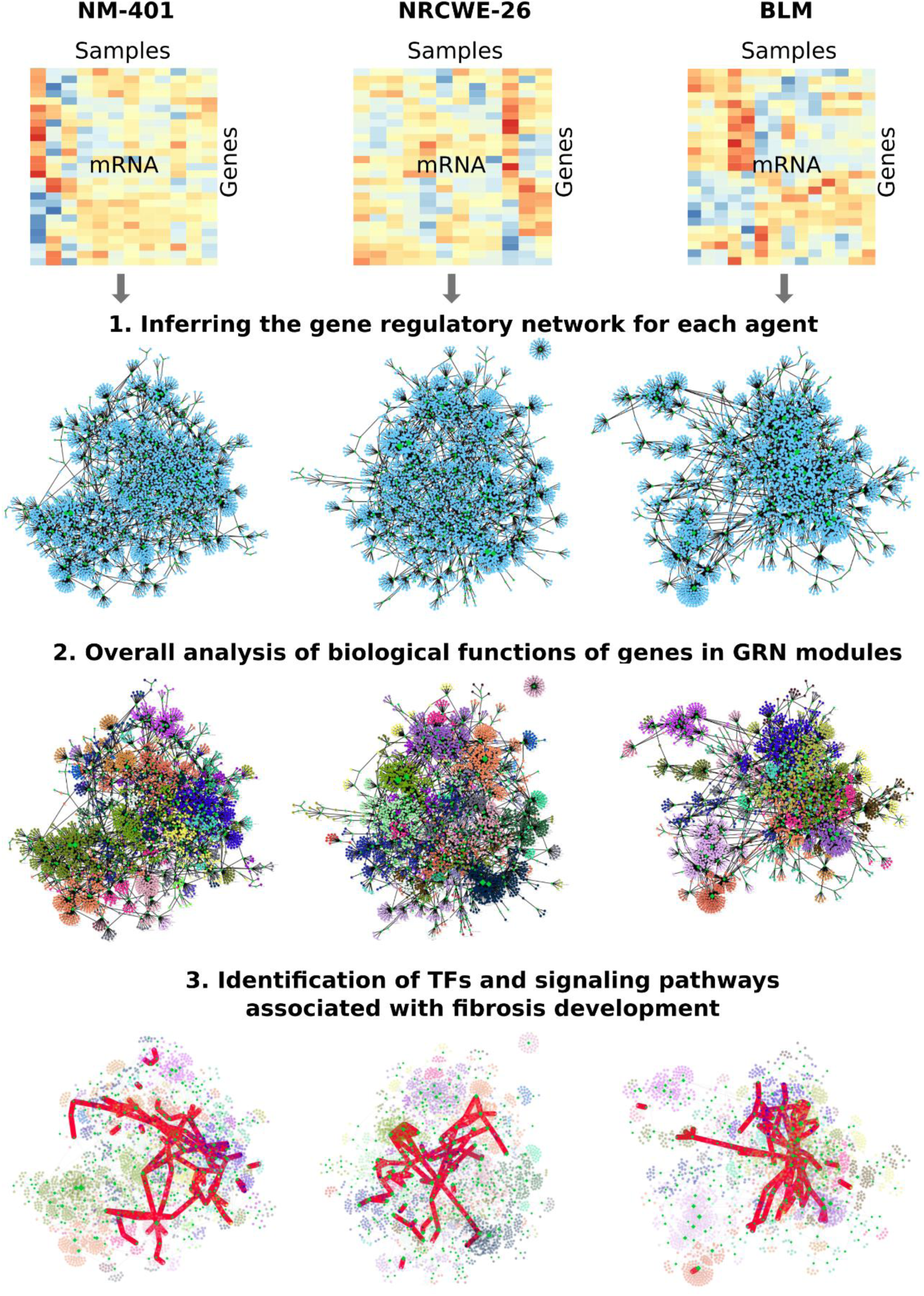
Workflow schematics for this study. To infer gene interactions from measured transcriptomics data, reverse engineering algorithms were applied. Using the clustering method, network modules were identified. For each module, pathway enrichment analysis was performed to identify the biological functions of the genes in the module. To reveal fibrosis-associated TFs and signaling pathways, subnetworks based on fibrosis markers and their direct regulators extracted from the whole networks were analyzed. The subnetworks are represented by bold red lines.

### 1. Inferring gene regulatory networks

Gene regulatory networks (GRNs) can provide useful insights into transcriptional regulatory mechanisms. GRNs inferred from the expression data can suggest which transcription factors (TFs) are responsible for the changes in gene expression observed following the exposure to two types of nanoparticles (NM-401 and NRCWE-26) and BLM. GRNs have hierarchical structures where a few highly interconnected genes, usually TFs, are the hubs that account for most interactions. It was previously shown that combining multiple inference methods increases the accuracy of reconstructed GRNs [15]. Therefore, we apply three different algorithms in order to infer GRNs: 1) linear model of gene regulation with Bayesian variable selection [16]; 2) mutual information algorithm ARACNe-AP [17]; and 3) random forest based algorithm GENIE3 [18]. To improve the prediction accuracy, we integrated the results of all three algorithms using the Borda count ranking method [15]. The list of TFs was used from AnimalTFDB database [19], and target genes were selected based on the combined list of DEGs from CNTs and BLM experiments. Analysis of DEGs was performed using the limma package in R/Bioconductor [20]. Genes were considered significantly differentially expressed if they (i) showed expression changes of at least ± 1.5-fold for CNTs or BLM treated groups compared to non-treated controls for each experimental condition; and (ii) have FDR p-values ≤ 0.05. The inferred networks characteristics are generally similar with clustering coefficients indicating that all GRNs are well-connected, small world networks (Table 1). However, the NM-401 GRN has a much larger diameter and a characteristic path length, suggesting that the activated regulatory program in response to NM-401 treatment might be more complex. The visualization of the networks is shown in Figure 2, and the cytoscape file can be found in the Additional file 1.

**Table 1.** Networks characteristics. The analysis was performed using the NetworkAnalyzer plugin available within the Cytoscape software.

| Network parameter | NM-401 | NRCWE-26 | BLM |
| --- | --- | --- | --- |
| Nodes | 3495 | 3559 | 3224 |
| Edges | 4802 | 4912 | 4438 |
| Transcription factors | 348 | 350 | 290 |
| Clustering coefficients | 0.058 | 0.05 | 0.048 |
| Network diameter | 16 | 8 | 9 |
| Shortest paths | 19549 | 12121 | 13807 |
| Characteristic path length | 3.295 | 2.146 | 2.384 |
| Avg. number of neighbors | 2.709 | 2.760 | 2.725 |

Next, in order to prioritize gene regulators, we identified TFs with the largest numbers of connections for each network, which will subsequently be referred to as TFs hubs. These network topological features are widely used in the analysis of GRNs and such types of genes are considered important in the cellular regulatory program [21, 22]. TFs were ranked based on their connection numbers (Additional file 2). The results show the TFs that have high connectivity in all networks, such as *E2f8* and *Irf7*. The other key TF hubs are differentially represented in different networks, such as Myc, *Ubtf, Etv4, Tbx20, Mafb* (Additional file 2).

### 2. Analysis of biological functions of genes in GRN modules

One of the known properties of GRNs structure is colocalization of genes from the same biological processes in the same network cluster [23]. We used this feature for a subsequent analysis of biological processes controlled by a TF. The inferred networks displayed modular structures (Figure 2, Additional file 1). To identify modules from these networks, the GLay clustering method in Cytoscape was applied [14]. Visualization of the clustered networks with upregulated and downregulated DEGs for different time points is presented in Additional file 3.

**Figure 2.**
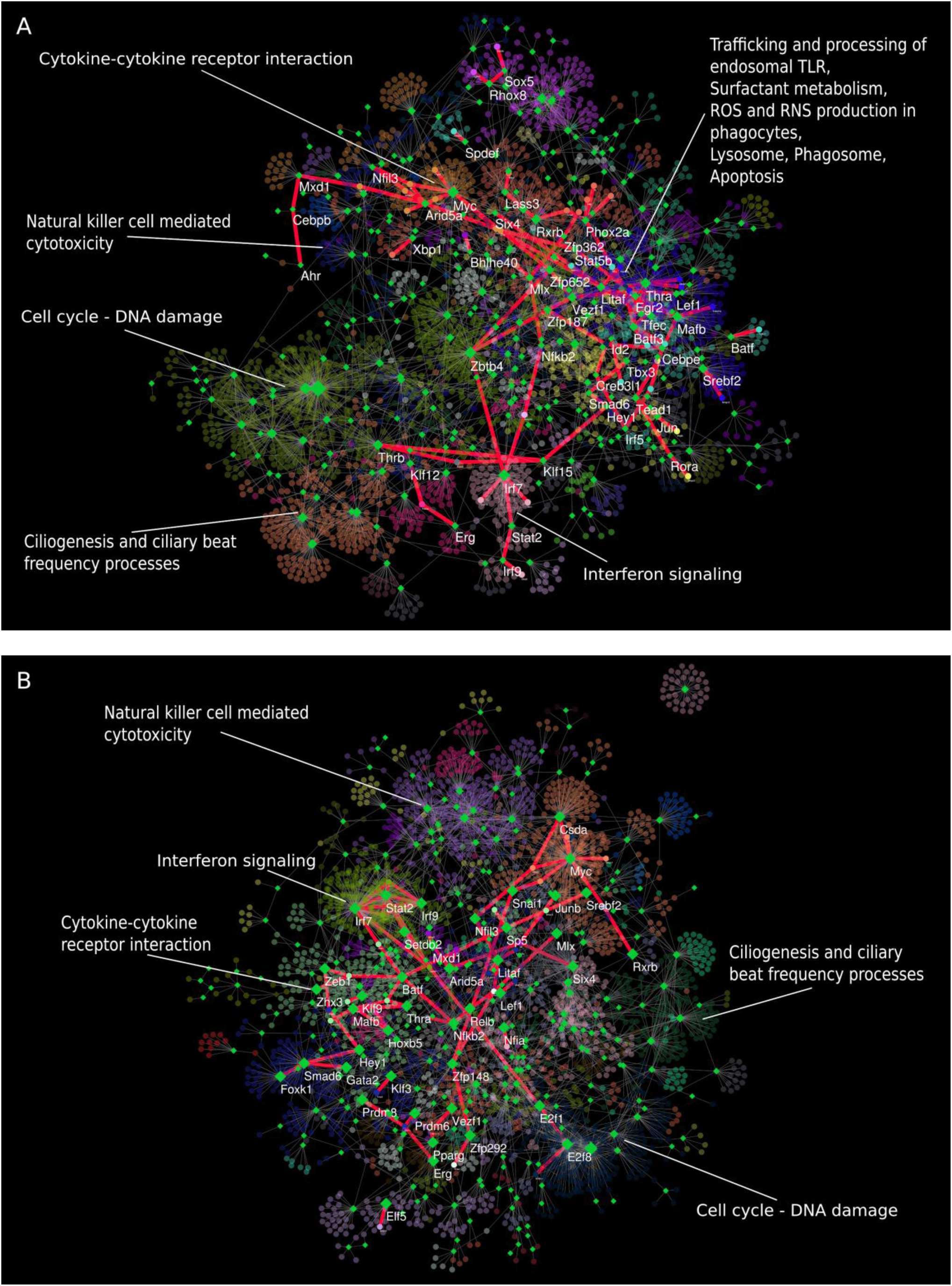
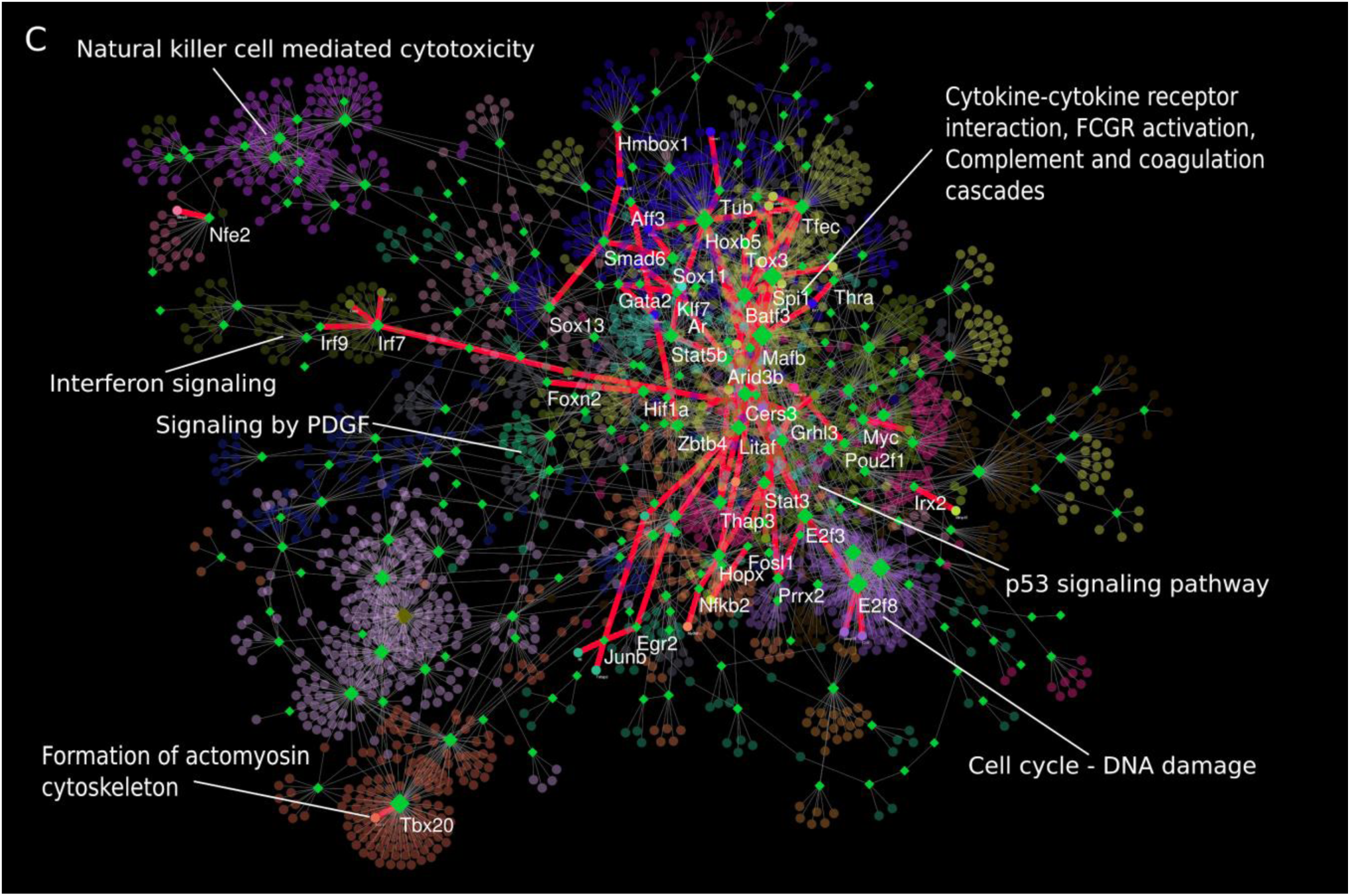
Inferred gene regulatory networks for carbon nanotubes and bleomycin. A) NM-401, B) NRCWE-26, C) BLM treatments. Green diamonds represent TFs, colored spherical nodes represent their potential targets and each color of spherical nodes indicate distinct module. The size of the diamonds is proportional to the number of external connections (outdegree). KEGG, REACTOME and GO databases were used for functional annotation of altered genes in the modules. Subnetworks based on fibrosis-related markers and their regulators are represented by bold red lines.

To identify the signaling pathways and functional processes which were altered in each module of the networks following instillation of CNTs and BLM, over-representation analysis was performed using gProfileR and ReactomePA toolkits [24, 25] and the KEGG, REACTOME and GO databases (see Additional files 4, 5, 6). The functional annotation analysis was performed for each time point and for upregulated/downregulated genes separately where the altered pathways had an adjusted p-value < 0.005. Instillation of CNTs and BLM affected various physiological and pathological mechanisms, such as the immunomodulatory response (innate and adaptive), response to DNA damage/integrity, pathways involved in cell-cell interactions and cell adhesion, and activation of regeneration processes, suggesting an involvement of these processes in the adverse effects observed upon CNTs and BLM instillation. We mainly focused on inflammation and fibrosis-related pathways, with the findings summarized in Table 2 and Additional file 7.

**Table 2.**
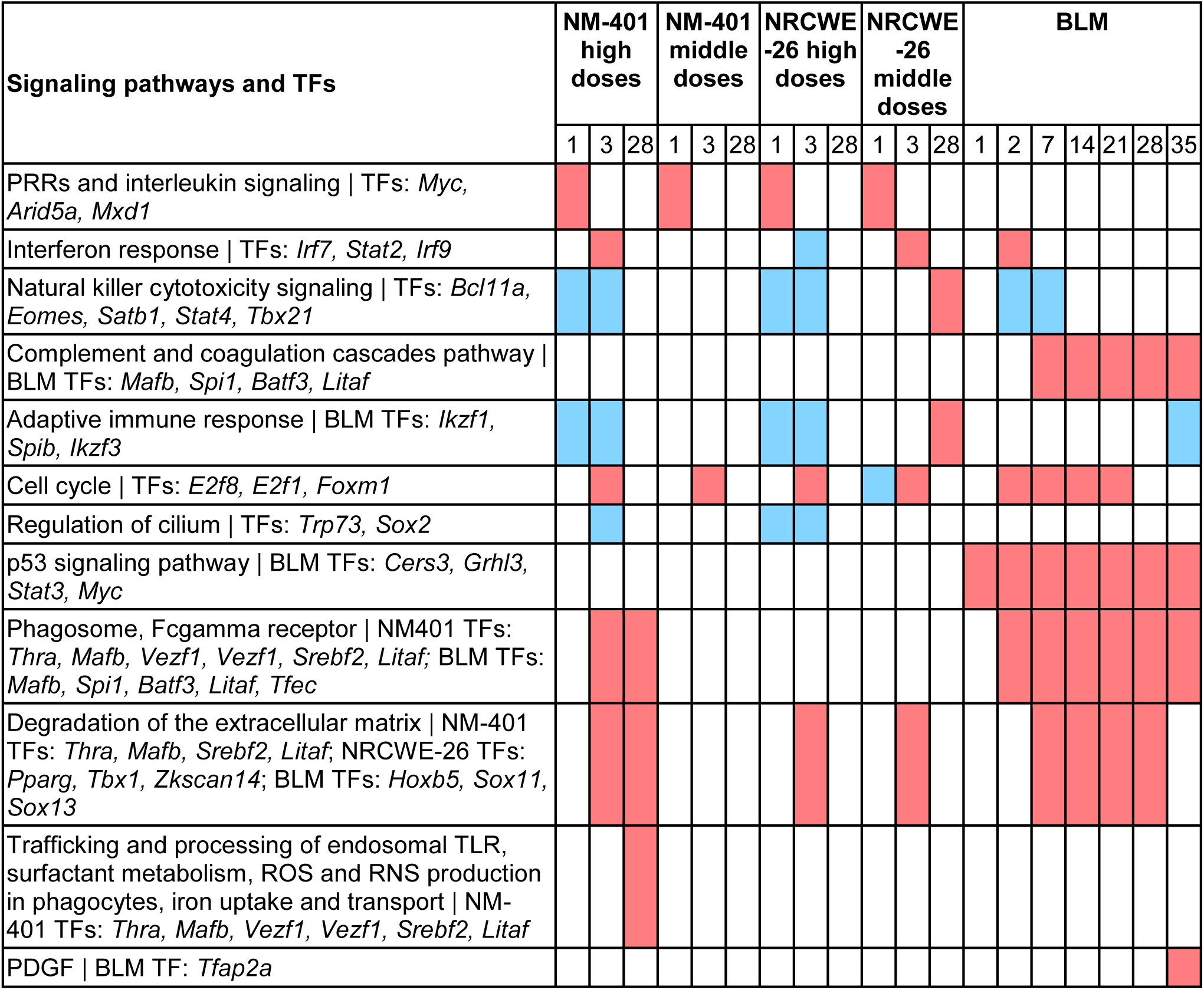
Summary of functional annotation analysis of GRN modules and associated TFs to identify pathways affected be CNTs and BLM exposure. The analysis was performed for each time point and for upregulated/downregulated genes separately. The adjusted p-value < 0.005 was taken as identifying the altered pathways and biological processes, which are shown by red and blue color for upregulated and downregulated genes, respectively.

The innate immune response was strongly stimulated by CNTs and BLM instillation. The components of pattern recognition receptors (PRRs) including Toll-like receptor, NOD-like receptor, RIG-I-like receptor, cytosolic DNA-sensing signaling pathways were upregulated in response to CNTs and BLM 1-3 days post-exposure. These pathways are typically activated in response to pathogen invasion and lead to the expression of cytokines and chemokines. The functional annotation analysis identified two types of modules in the networks, which were associated with alternative downstream signaling of PRRs. DEGs from the first type of module generally were upregulated 1-2 days post-exposure and belonged to PRRs pathways and interleukin signaling, while DEGs from the second type of module were activated on 2-3 days and belonged to PRRs and Interferon signaling pathways. Both the high dose of NM-401 and the middle dose of NRCWE-26 induced an upregulation of DEGs in the Interferon module, while BLM induced a moderate effect. Interestingly, the high dose of NRCWE-26 inhibited the expression of DEGs from this particular module (see Figure 3, Additional file 7). Our analyses revealed common TF hubs in modules with similar functions in the different networks; for example, *Arid5a, Mxd1* were identified as TF hubs for the PRR-Interleukin module, while *Irf7, Stat2* and *Irf9* were identified as TF hubs for PRR-Interferon module. In addition, other innate immune response pathways also were altered (Additional files 4, 5, 6, 7). All of the assessed agents were shown to affect signaling associated with natural killer cytotoxicity. DEGs involved in this signaling were downregulated in response to BLM on 2-7 days and high doses of CNTs on 1-3 days, and upregulated in response to middle doses of NRCWE-26 on 28 day. The TFs *Bcl11a, Eomes, Satb1, Stat4, Tbx21* were identified in the natural killer cytotoxicity module in all networks. BLM also affected the complement and coagulation cascades pathway with DEGs upregulated at 7-35 days. Components of adaptive immune response, such as B cell signaling, were downregulated in response to BLM and high doses of NRCWE-26. Likewise, DEGs from Th1/Th2 cell differentiation, antigen processing and presentation pathways were downregulated in response to high doses of NRCWE-26.

The expression changes of the innate immune system components were followed by changes in the cell cycle and DNA damage signaling modules. DEGs in these modules were upregulated in response to all agents on day 3 (CNTs) / 2-21 (BLM), and downregulated on day 1 in response to middle dose of NRCWE-26. These DEGs contained genes from homologous recombination, DNA replication, cell cycle checkpoints, p53 and other signaling pathways which are involved in cell cycle and DNA damage processes. The TF hubs identified in this module were *E2f8, E2f1, Foxm1*.

The cell cycle module has a high number of connections with a closely located module containing the TF hubs *Trp73* and *Sox2* in the CNTs networks (Figure 2). DEGs from the module were downregulated in response to NRCWE-26 on day 1, and NM-401 and NRCWE-26 on day 3 (see Additional file 3). Functional analysis of these genes revealed a relationship with ciliogenesis and ciliary beat frequency processes (GO BP ‘axoneme assembly’, ‘regulation of cilium beat frequency’). Cilia are cell organelles in lung epithelial cells which provide airways clearance from mucus and dirt. Attenuation of ciliogenesis in response to CNTs has been shown previously in bronchial epithelial cells [26].

All doses of NRCWE-26 inhibited DEGs from the cardiac muscle contraction pathway on day 3, while in the case of BLM this inhibition began at day 14 and was maintained even after day 28. The identified genes belonged to the actin and myosin family of genes, which are involved in the formation of the actomyosin cytoskeleton of non-muscle cells. Disruption of this structure in phagosome cells can facilitate engulfment processes [27]. TF *Tbx20* was identified as TF hub in this module.

Other signaling pathways which were altered during the later time period included phagosome and osteoclast differentiation (KEGG). DEGs from these pathways were shown to be upregulated from the earliest time period and were continuously activated up to the 28^th^ day (NM-401) and 35^th^ day (BLM). Based on gene ontology analysis, the enriched genes from these pathways belong to the Fcgamma receptor (FCGR) family, ATPases and macrophage scavenger receptors. Enrichment analysis using the REACTOME database also revealed alteration of the FCGR signaling pathway. It is conceivable that treatments activated FCGR-dependent phagocytosis processes. The FCGR signaling is activated by immunoglobulin G (IgG) molecules [27] and plasma protein serum amyloid P (SAP) [28]. IgG antibodies can be contained in the protein corona of nanoparticles and act as opsonins that by binding to phagocyte FCGRs initiate the re-organization of the actomyosin cytoskeleton, membrane remodeling, and eventually phagocytosis [27]. Furthermore, the FCGR can be activated by SAP protein during clearance of apoptotic cells [28]. Likewise, the same module of BLM network consisted of upregulated DEGs (from day 7) from the complement and coagulation cascades pathway, which also can enhance opsonization. This upregulation can be explained by the activation of apoptotic material clearance processes. All three pathways, phagosome, osteoclast differentiation and coagulation cascades, were identified in the same modules, which consisted of the TF hubs *Mafb, Spi1, Batf3, Litaf* and *Tfec* in the BLM network and TF hubs *Thra, Mafb, Vezf1, Vezf1, Srebf2* and *Litaf* in the NM-401 network.

Several other pathways altered by CNTs and BLM included the following. DEGs mapped to the degradation of the extracellular matrix pathway were upregulated in response to all treatments, but only BLM and NM-401 affected this process at the later time periods (days 7-28 BLM, and day 28 NM-401). Pathways altered in response to NM-401 at day 28 included the trafficking and processing of endosomal TLR, surfactant metabolism, ROS and RNS production in phagocytes, iron uptake and transport pathways. Components of all these different pathways were upregulated and were enriched in the module with the TF hubs *Thra, Mafb, Vezf1, Vezf1, Srebf2* and *Litaf*. PDGF signaling was altered in response to BLM, with the DEGs within this pathway upregulated at day 35. PDGF activity can be essential for myofibroblast activation, which is one of the important players in fibrosis development. *Tfap2a* was identified as TF hub gene in this module. The late time period effects of NRCWE-26 depended on its dose. High NRCWE-26 dose, as well as BLM, induced upregulation of inflammatory processes. DEGs from NOD-like receptor and cytokine-cytokine receptor interaction pathways were activated. Middle dose NRCWE-26 induced adaptive immune system processes, such as autoimmune thyroid disease and antigen processing-cross presentation. Furthermore, components of natural killer cell mediated cytotoxicity and cellular senescence were upregulated.

**Figure 3.**
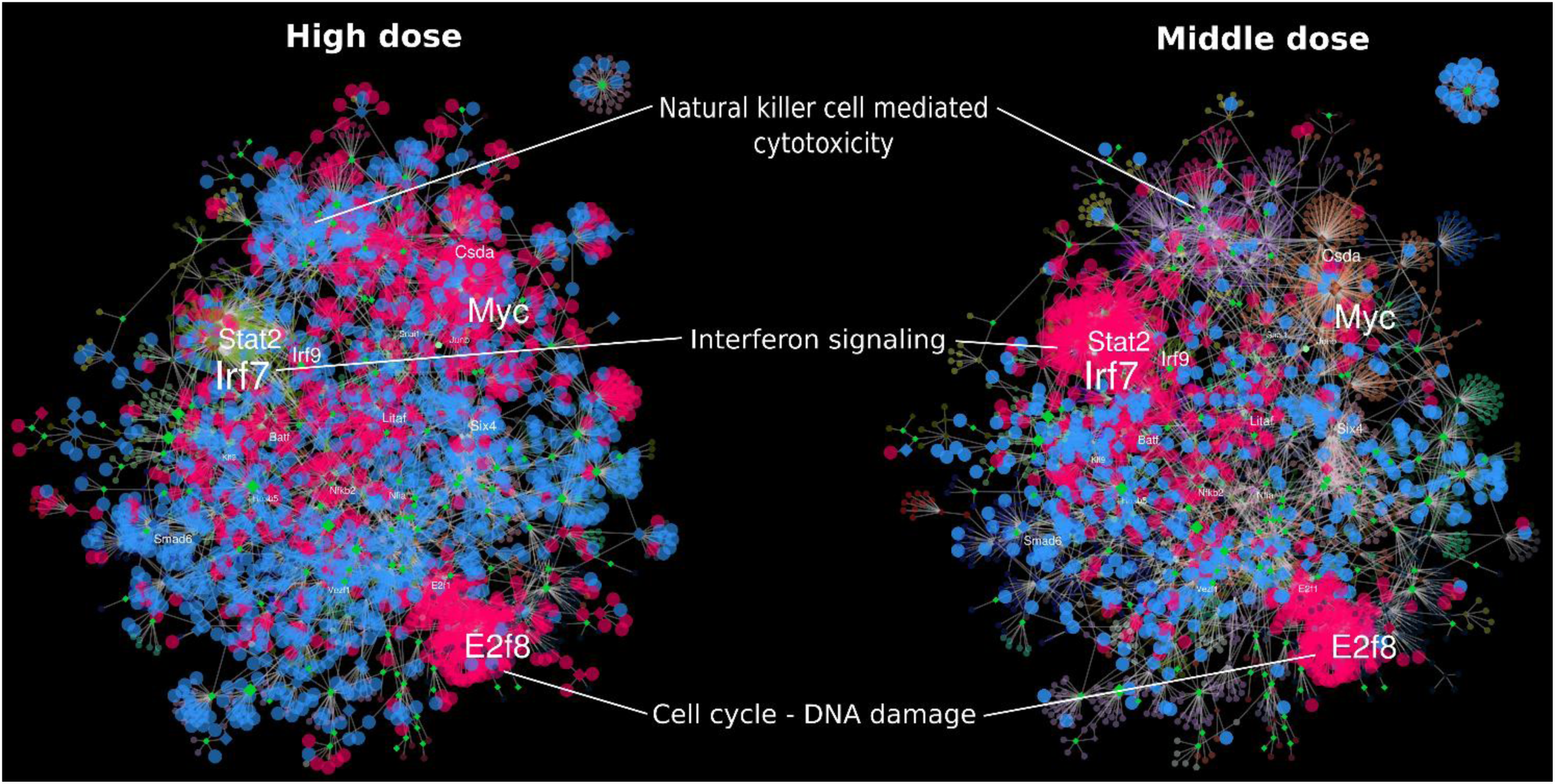
Visualization of DEGs in the NRCWE-26 GRN. The red and blue colors represent the upregulated and downregulated DEGs, respectively, for high doses (on the left) and middle doses (on the right) at the day 3 timepoint.

Taken together, our results indicate that instillation of these CNTs and BLM activated various physiological and pathological mechanisms, such as inflammation, response to DNA damage, alteration in ECM synthesis and activation of the regeneration process, thus highlighting the involvement of these processes in the adverse effects observed upon CNTs and BLM instillation. As indicated by the data for later time period responses, NM-401, as well as BLM, led to remodeling of the extracellular matrix and active phagocytosis. Interestingly, NRCWE-26 induced biphasic effects: high doses of NRCWE-26 activated the innate immune response, while middle doses triggered the adaptive immune components. Moreover, middle doses of NRCWE-26 induced the interferon response, while high doses did not alter the genes from this module (see Figure 3). Heatmap analysis of DEGs from the Interferon signaling pathway (where the list of genes was derived from REACTOME Interferon signaling pathway, R-MMU-913531) revealed the same effect for NRCWE-26 exposure (Additional file 8).

### 3. Analysis of gene regulators

In the previous section, we identified main TF hubs for functionally annotated modules. These gene regulators were associated with the following different responses: *Arid5a, Mxd1* for interleukin signaling; *Irf7* for interferon response; *E2f8, E2f1, Foxm1* for cell cycle and DNA damage response; *Trp73* for cilium regulation; *Mafb* for phagocytosis and Fcgamma receptors activation; *Tbx20* for actomyosin cytoskeleton remodeling. It was, however, unclear, whether these regulators were important for the main pathological effect of CNTs and BLM treatment such as pro-fibrotic responses.

To gain more insight into the role of gene regulators during the activation of fibrotic processes, we next focused on specific subnetworks within the GRNs. Results were obtained by using 87 genes, which were previously identified to be important for fibrogenesis and tissue remodeling in response to different types of CNTs [13, 29]. This gene set includes several matrix metallopeptidases and their inhibitors, interleukins, chemokines, ECM regulators, and other genes which are involved in parenchymal injury of the lung. We considered this list of genes as targets and found their regulators using the inferred whole GRNs for NM-401, NRCWE-26 and BLM (as mentioned above). These genes and their regulators formed a unique subnetwork and linked different functional modules in the whole network (Figure 2). Previously identified functional modules, such as cell cycle - DNA damage, interleukins, interferon and phagocytosis were interlinked by this subnetwork, suggesting that they contribute to the fibrotic response *via* a coordinated transcriptional program.

To identify which TFs are prevalently involved in the regulation of DEGs at different time points, we reconstructed subnetworks based on DEGs from the gene set for each time point and found their regulators in the whole GRNs. The following analysis was performed based on the identified number of connections for each TFs (Table 3). Myc, *Arid5a* and *Mxd1* had a high number of connections in CNT subnetworks at days 1-3. *Arid5a* and *Mxd1* were included in PRR-interleukin module, which was activated in response to CNTs treatment at day 1-3 (as mentioned above). *Myc* was identified in the top 3 TF hubs with the highest number of connections in all CNT networks (See section “Inferring gene regulatory networks”, Additional file 2), indicating that this TF plays an important role in cell regulation in response to CNT treatment. In BLM network Myc was associated with p53 signaling (Table 2, Additional file 7). *Myc* activity is essential for cell cycle progression, apoptosis and other biological processes. Myc has cross-regulatory interactions with cytokines, including Il1, Il2, Il4, Il6, Il8, Il10, and TNF-α [30]. In NM-401 GRN *Myc* colocalized with *Arid5a*, which had also the high number of connections in the subnetworks (see Table 3). The TF *Arid5a* has a role in the posttranscriptional regulation of IL6. *Arid5a* controls *Il6* mRNA stability and protects *Il6* mRNA from regnase-1-mediated degradation [31]. Importantly, *Arid5a* is regulated by the NF-κB and MAPK signaling pathways, which in turn are activated by Toll-like receptor 4 [32]. Another TF with high number of connections in CNT subnetworks was *Mxd1*, also known as *Mad*. This TF is involved in MYC regulation.

**Table 3.**
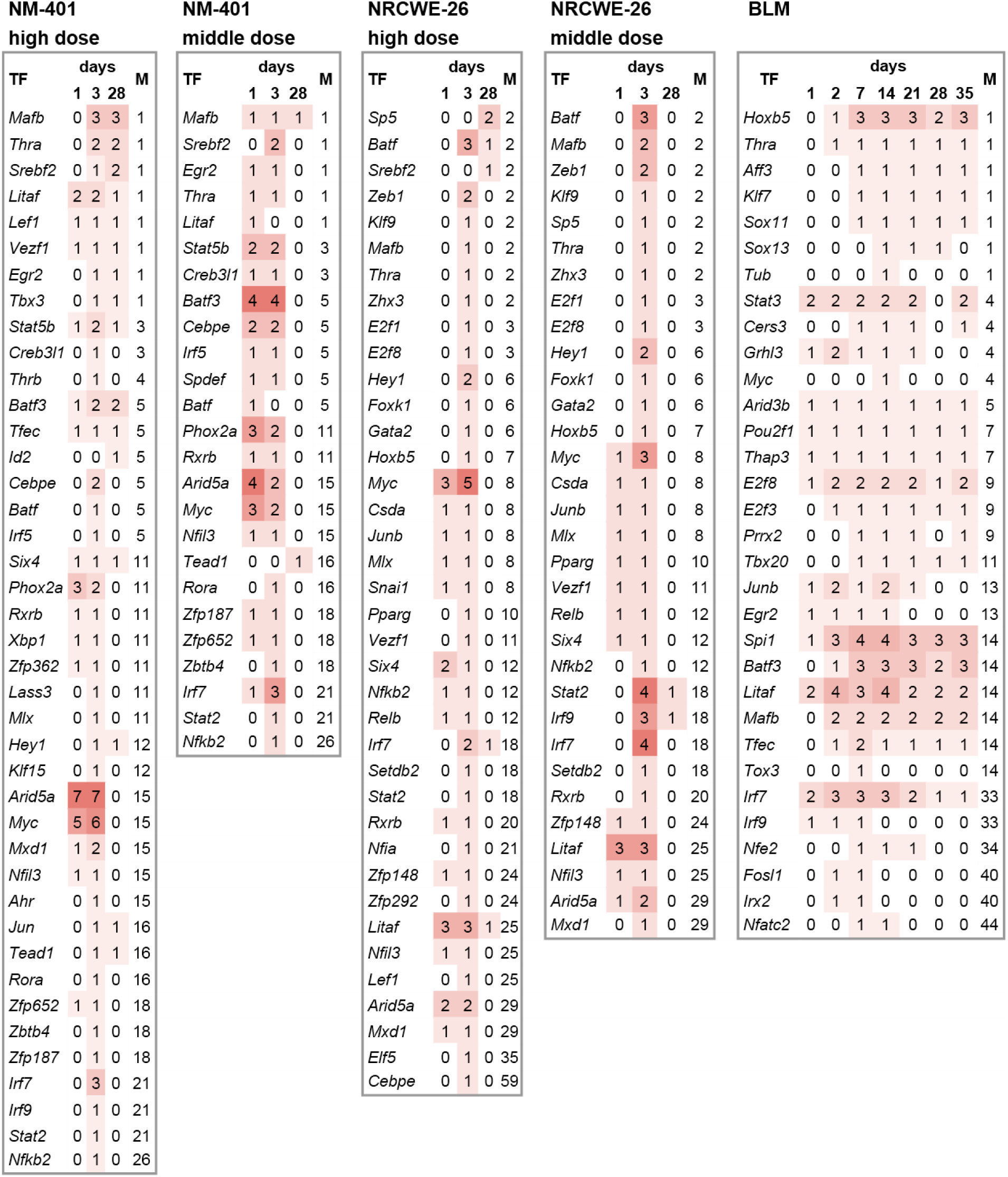
Connectivity patterns for TFs. The number of connections for each TF is shown. M represents the number of modules in the corresponding subnetwork.

Heterodimerization of MYC with MAX is necessary for activation of MYC target genes. The protein MAD, which is encoded by *Mxd1* gene, competes with MYC for binding to MAX and thereby inhibits MYC activity [33]. Analysis of the log fold change (logFC) values for *Myc, Max, Mxd1* and cytokines showed that the largest *Myc* to *Mxd1* ratio (ΔlogFC 1.66) was observed for day 3 for high dose of NRCWE-26 (Additional file 9).

Other TFs with a high number of connections in all subnetworks were identified. *Litaf* regulates the expression of cytokines, pro-inflammatory and pro-fibrogenic genes [34–36]. Transcription of *Litaf* can be induced by tumor suppressor p53 [37] and Toll-Like receptors (TLR 2/4) [38]. In the CNT networks *Litaf* is directly connected with *Cd14* and *Myd88* genes (see Additional file 10), which encode toll-like receptor interacting proteins. The TF *Irf7* from the PRR-interferon GRN module (as mentioned above) also had high degree of connectivity in all subnetworks, especially at early time points (Table 3). This TF plays essential role in the activation of the viral defense system *via* triggering type I interferon pathway [39]. TFs *Mafb* and *Batf3* had high connectivity degree in NM-401 and BLM subnetworks at late time points (Table 3). *Mafb* was identified in a module associated with FCGR activation and phagocytosis (as mentioned above), which was altered also at late time points in the NM-401 and BLM networks. *Mafb* can enhance phagocytic activity of macrophages by stimulating Fcgr3 [40] and has a key role in the activation of anti-inflammatory profile of macrophages by inducing M1/M2 macrophage polarization, which is important for fibrosis development [41, 42]. Macrophages produce numerous cytokines, chemokines, matrix metalloproteinases, and other inflammatory and ECM remodeling mediators. Macrophages can be transformed by external stimuli into different types: M1 (pro-inflammatory) and M2 (anti-inflammatory) subtypes. The other TF, *Batf3*, is involved in formation of CD103+ and CD8+ dendritic cells that may facilitate lung fibrosis [43, 44]. In line with this result, liver fibrosis was attenuated in *Batf3^−/−^* knockout mice [45]. The TF *Srebf2*, also known as *Srebp2*, was increasing its connectivity over time in the CNT subnetworks for high doses of these agents. *Srebf2* induces the expression genes that are involved in cholesterol and fatty acid synthesis, cholesterol transport [46, 47] and in the formation of lipid-laden macrophages (foam cell) [48], which are associated with lung fibrosis [49].

Surprisingly, TFs from the cell cycle and DNA damage module, which included dozens of upregulated genes in all networks (as mentioned above), were weakly represented in the CNTs subnetworks. By contrast, analysis of the BLM subnetwork showed a high number of connections for these TFs. For instance, in the BLM network *E2f8, E2f3* and *Prrx2* had high degrees of connectivity up to late time points. Thus, the activity of genes from Cell cycle and DNA damage module was mainly associated with fibrotic changes in response to BLM.

## Discussion

Transcription factors regulate stress-induced biological processes and determine the landscape of toxicological response [50–52]. Characterization of the gene regulatory programs activated in response to nanoparticle treatment can improve risk assessment and potentially aid to build predictive toxicity models. Here, we applied reverse engineering techniques for reconstructing gene regulatory networks based on microarray mRNA data from the lungs of mice exposed to MWCNTs (NM-401, NRCWE-26) and BLM. Previous studies have demonstrated the power of this approach to identify molecular biomarkers and drug targets in cancer and other diseases [53], the deduction of adverse outcome pathways (AOPs) for chemicals from high-throughput transcriptomic data sets [54, 55], and the functional interpretation of responsive modules from gene expression data sets [56]. Additionally, based on prior knowledge of interactions between genes and a predefined list of disease-associated markers, the pipeline for detecting AOP-linked molecular pathway descriptions has been described [57].

Since TF activity is dependent on coactivators and because a single protein can be involved in different biological processes, as the first step, we identified functional roles of TFs using so-called “guilt by association” approach [23, 58]. The inferred whole networks were clustered and functional enrichment analysis was performed for each gene module. Additional analysis applied to the whole networks is a deduction of gene regulators associated with previously identified fibrosis markers. For this purpose, we inferred a subnetwork based on 87 genes (which were previously identified as important for fibrogenesis and tissue remodeling in response to different types of CNTs [13, 29]) and their direct regulators identified in the whole networks (see Figure 2).

*Arid5a, Myc* and *Mxd1* were identified as important gene regulators of the acute inflammatory phase under stress conditions developed following CNT treatment. *Arid5a* was identified as a TF hub in the interleukin module in the whole CNT networks and fibrosis-related subnetworks. This TF was directly connected with *Il6* gene in CNT networks, suggesting their closely related activity. A few studies have been published that demonstrate that *Arid5a* is an essential player in the posttranscriptional regulation of Il6. *Arid5a* controls *Il6* mRNA stability and protects *Il6* mRNA from regnase-1-mediated degradation [31]. Importantly, *Arid5a*-deficient mice display reduction of BLM-induced lung injury [59]. *Arid5a*, together with the TF *Myc*, colocalized in the CNT networks with cytokine-related genes, mostly appearing in the first phase of inflammatory response. *Myc* also was identified as a TF hub in whole CNT networks and *Myc* had one of the highest number of connections in the fibrosis-related subnetwork. This finding is in line with a recent report that upstream analysis identified *Myc* as a key regulatory gene, activated in response to both NM-401 and NRCWE-26 treatments [6]. *Myc* may also be involved in fibrogenetic processes. Myc can regulate and be itself regulated by several cytokines [30], which stimulate the expression of pro-fibrotic genes. On the other hand, among Myc’s transcriptional targets are integrins, which are transmembrane, cell signaling proteins that control cell surface adhesion, cell-matrix and other processes [60, 61]. Integrins can mediate the activation of the latent form of TGF-β [61–63], which is considered as a key regulator of fibrosis. *Myc* can directly bind to the promoter of integrin αv and induce its transcription [61], thereby promoting the activation of the latent form of TGF-β. Activation of *Myc* target genes depends on its cofactors. Heterodimerization of Myc with Max is necessary for activation of Myc target genes. The protein Mad, which is encoded by *Mxd1* gene, competes with Myc for binding to Max and thereby inhibits Myc activity [33]. In our analysis, *Mxd1* was also identified as a crucial TF. Therefore, we hypothesize that the balance between the fold change value of *Myc* and *Mxd1* can be used for the prediction of *Myc* activity. The ratio *Myc : Mxd1* was significantly higher in NRCWE-26 than in NM-401-induced stress response (1.66 and 0.3, respectively). This result corroborates the finding that the logFC for *Myc* is greater for NRCWE-26 than for the NM-401 treatment (Additional file 9).

Transcription factor *Irf7*, the important player in the innate immune response and the activator of the viral defense system *via* triggering type I interferon pathway [39], was identified in this analysis as a critical gene regulator in response to both CNTs and BLM. The most interesting finding was that NRCWE-26 inhalation showed a biphasic effect: middle and low doses of NRCWE-26 induced the interferon response, while high doses did not alter genes from this signaling pathway. A possible explanation for this effect may be that in high concentrations NRCWE-26 nanoparticles can form agglomerates (clots), which are sensed by immune cells in a different manner than distinct nanoparticles.

Another transcription factor that has been identified as an inflammation-related gene regulator for CNTs and BLM treatment is *Litaf*, which controls one of the alternative downstream signaling pathways of TLR2/4 [38]. These types of toll-like receptors control interferons and cytokines expression and are involved in cellular stress response upon action of exogenous or endogenous ligands, such as bacterial LPS and group of proteins from damage-associated molecular pattern [64]. Data from several sources have identified the activation of TLR2/4 signaling in response to nanoparticles [65, 66] or BLM treatment [67]. *Litaf* was upregulated during acute inflammation phase and had direct links with fibrosis-related genes in CNT and BLM networks for the later time period.

Our findings further support the idea of a pivotal role of macrophage polarization in the cellular response to nanoparticles and BLM [12, 68]. We identified the TF *Mafb* as a potential driver of this activity. *Mafb* was identified in the networks associated with Fc gamma receptors expression (as mentioned above), which can define pro- or anti-inflammatory profile of immune cells [69] and also can be involved in regulation of M1/M2 macrophage polarization [70].

We have also shown that cell cycle and DNA damage signaling was altered in response to all agents. TFs *E2f8, E2f1, Foxm1* were identified as important regulators of this stress response signaling. It is interesting to note that cell cycle and DNA damage module was mainly associated with fibrotic markers in the BLM network. These findings further support the idea of different nature of activation of cell cycle and DNA damage pathways in response to BLM and nanoparticles. BLM can directly induce single- and double-stranded DNA breaks [71], while NM-401 and NRCWE-26 nanoparticles induce mainly an inflammation response, which can cause DNA damage. Previous studies have reported the absence of DNA strand breaks for these nanoparticles [11, 13].

## Conclusions

In the present study we revealed the landscape of transcriptional regulation of responses to CNTs and BLM. This analysis identified common and distinctive features for each agent. Activity of *Myc* was mainly associated with cytokine regulation in response to nanoparticles and with p53 signaling in response to BLM. Under the control of TFs *Irf7, Stat2* and *Irf9*, interferon response was activated by all agents, and NRCWE-26 treatment showed a biphasic, dose-dependent effect. Despite the different nature of the lung injury caused by CNTs and BLM, we identified common gene regulators. TFs *Litaf* and *Mafb* were identified as essential regulators in noncanonical signaling of TLR2/4 and M1/M2 macrophage polarization, respectively, highlighting the universal features of these cell responses for fibrosis progression.

In summary, we developed the TF-centric pipeline used to reveal gene regulators, their associated biological processes and signaling pathways, which were altered in response to CNTs and BLM. This method uses transcriptomics data, generates specific to toxic agent interaction networks and is independent from bias in the reference databases for pathway mapping. Moreover, this approach can be useful for generating toxicity pathways and adverse outcome pathway schemes for toxic agents.

## Methods

### Data set

The gene expression profiles for two types of multi-walled CNTs (GSE55286), NM-401 (4048±366 nm in length) and NRCWE-26 (847±102 nm in length), and bleomycin (GSE40151) were obtained from Gene Expression Omnibus Database (https://www.ncbi.nlm.nih.gov/geo/) (Table 4). Data for the MWCNTs was generated using Agilent SurePrint G3 Mouse GE 8×60K Microarrays and for bleomycin using Affymetrix Mouse 430 2.0 arrays in *in vivo* experiments. In the nanoparticle experiments three different doses of CNTs (18, 54, 162 μg) and three post-instillation timepoints (1, 3, 28 days) were employed. Bleomycin was administered with the one dose 2U/kg body weight, the lung tissue was harvested at 7 post-instillation timepoints (1, 2, 7, 14, 21, 28, 35 days). Both type of experiments were conducted with vehicle controls for each timepoint.

**Table 4.** List of used data sets from Gene Expression Omnibus Database.

| Data set | Publication | Treatment | Mouse strain | Number of samples | Post-exposure time point, days | Doses | Type of microarray |
| --- | --- | --- | --- | --- | --- | --- | --- |
| GSE55286 | [13] | MWCNT (long NM-401 4048±366 nm, short NRCWE-26 (847±102 nm)) | C57BL6 | 70 (NM-401), 69 (NRCWE-26) | 1, 3, 28 | 18, 54, 162 µg, vehicle control | Agilent SurePrint G3 Mouse GE 8x60K Microarray |
| GSE40151 | [9] | Bleomycin | C57BL6 | 111 | 1, 2, 7, 14, 21, 28, 35 | 2U/kg, vehicle control | Affymetrix Mouse 430 2.0 arrays |

DEG analysis was performed using the limma package in R/Bioconductor [20]. The list of genes was considered as significantly differentially expressed if the expression changes were equal to or larger than ± 1.5-fold for nanoparticles or bleomycin treated group compared to non-treated controls for each experimental conditions and the BH-adjusted (FDR) p-values were less than or equal to 0.05 (p ≤ 0.05).

### Gene Regulatory Network inference

In order to infer gene regulatory networks for two types of MWCNTs and BLM, we applied three different algorithms: linear model of gene regulation with Bayesian variable selection [16], the mutual information algorithm ARACNe-AP [17], and the random forest based algorithm GENIE3 [18]. The gene interactions were identified for the combined list of DEGs derived from all experiments. The predefined list of gene regulators (TFs) was used from the AnimalTFDB database [19]. Next, for improving prediction accuracy, we integrated the results of all three algorithms by Borda count ranking as described previously [15]. The ARACNe-AP algorithm was run with three key steps: MI threshold estimation, bootstrapping/MI network reconstruction, building consensus network (only significant interactions are filtered, p < 0.05, Bonferroni corrected). Bayesian variable selection and GENIE3 algorithms were run with default parameters [16, 18]. Top 3% of the ranked edges in each common network were selected for subsequent analysis. Network visualization, parameter analysis were performed by open source software platform Cytoscape version 3.4.0 [72]. The integration of initial GRNs, data processing, statistical analysis were performed with R version 3.3.3 (https://www.r-project.org/) and RStudio version 1.0.44 (https://www.rstudio.com). To identify GRNs modules, the GLay clustering method in Cytoscape was applied [14]. For functional annotation of GRN modules we used KEGG [73], REACTOME [74], GO [75] databases and gProfileR, ReactomePA toolkits [24, 25]. A threshold for the minimum number of genes per module was 30 and BH-adjusted (FDR) ρ-value threshold was 0.005.

## List of abbreviations

BLM: : bleomycin
CNTs: : Carbon nanotubes
DEGs: : differentially expressed genes
ECM: : extracellular matrix
FCGR: : Fcgamma receptor
GRN: : gene regulatory network
MWCNT: : muli-walled carbon nanotubes
PRRs: : pattern recognition receptors
TF: : transcription factor

## Funding

This work was supported by the EU H2020 SmartNanoTox (Grant no. 686098).

## Additional files

Additional file 1: Inferred GRNs in cytoscape file format.

Additional file 2: The number of connections for the top 20 (the cells filled with green colour) of largest TFs hubs for each whole network.

Additional file 3: Visualization of the clustered networks with upregulated and downregulated DEGs for different time points.

Additional file 4: KEGG pathway analysis of differentially expressed genes from GRN modules. The analysis was performed for each time point and for upregulated/downregulated genes separately.

Additional file 5: REACTOME pathway analysis of differentially expressed genes from GRN modules. The analysis was performed for each time point and for upregulated/downregulated genes separately.

Additional file 6: GO functional analysis of differentially expressed genes from GRN modules. The analysis was performed for each time point and for upregulated/downregulated genes separately.

Additional file 7: Summary functional analysis of GRN modules.

Additional file 8: Heatmaps of DEGs from Interferon signaling pathway (REACTOME).

Additional file 9: Balance of *Myc* and *Mxd1*. The experimental results are presented in the descended order of logFC values for Myc. List of cytokines were derived from “Innate & Adaptive Immune Responses PCR Array” RT^2^ Profiler PCR Array System, QIAGEN.

Additional file 10: First neighbors for *Litaf*.

